# A Bioelectronic Scaffold for Label-Free, Real-Time Monitoring of Wound Healing

**DOI:** 10.1101/2025.09.04.674270

**Authors:** Dana Cohen-Gerassi, Offir Loboda, Aakash Jog, Or Messer, Dor Aaron Goldstein, Tomer Reuveni, Moran Aviv, Amit Sitt, Yosi Shacham-Diamand, Maxim Sokol, Lihi Adler-Abramovich

## Abstract

Chronic wounds and severe burns pose a major clinical challenge, as they often heal slowly or fail to respond to conventional treatments. In addition, there is a critical lack of tools for personalized, continuous monitoring of the healing process. Although progress has been made in both regenerative biomaterials and wearable biosensors, their integration into a unified platform that enables *in situ*, real-time monitoring of wound healing remains a major challenge. Here, we present a multifunctional bioelectronic scaffold that combines regenerative capability with real-time sensing of cellular activity. The scaffold was fabricated by electrospinning polycaprolactone (PCL) functionalized with the bioactive, self-assembling peptide fluorenylmethoxycarbonyl-phenylalanine-arginine-glycine-aspartic acid (Fmoc-FRGD) to promote cell adhesion and proliferation. For electrical sensing, biocompatible MXene (Ti_3_C_2_T_X_) electrodes were conformally deposited onto the nanofibrous matrix, preserving its biological functionality.

This system enables label-free, real-time monitoring of cell viability and coverage using electrical impedance spectroscopy (EIS), offering continuous and quantitative insight into cellular adhesion and proliferation. Extracted impedance parameters at low frequencies exhibit a strong correlation with both cell viability and coverage, providing a non-destructive indicator of wound closure and healing dynamics. This platform offers a promising strategy for advanced wound care, integrating real-time monitoring with biologically supportive materials.

## Introduction

Chronic wounds, including diabetic ulcers, pressure sores, and severe burn injuries, represent a growing global health burden, affecting up to 2 % of the population in developed countries and resulting in prolonged hospitalization, recurrent infections, and increased risk of limb amputation or mortality.^1–3^ Despite advances in regenerative medicine and wound care technologies, these wounds often fail to progress through the normal stages of healing and remain resistant to standard therapies. As a result, they impose a substantial socioeconomic burden, with the annual cost of chronic wound care in the United States alone estimated at $28-97 billion.^4^

The clinical management of chronic wounds and burns is further limited by the lack of objective and reliable tools for real-time assessment of healing progression. Today, clinicians rely primarily on periodic visual inspection during dressing changes to monitor wound status, a method that is inherently subjective, infrequent, and incapable of capturing early signs of infection or tissue remodeling at the cellular level. Moreover, each dressing change disrupts the wound microenvironment, potentially impeding the healing process, while also subjecting the patient to considerable pain and discomfort. In response to the clinical need for skin regeneration, several engineered skin substitutes and advanced wound therapies have been developed, including biological grafts, synthetic scaffolds, and commercial products such as Integra®^5^, Dermagraft®^6^, and Epicel®^7^. Among these, Cultured Epidermal Autografts (CEAs), composed of autologous keratinocyte sheets expanded *ex vivo*, remain a widely used approach for treating extensive burns.^8^ However, CEAs are fragile, costly, and difficult to handle, lacking both dermal components and the mechanical strength required for effective full-thickness skin regeneration.^9,10^

Peptide-functionalized scaffolds offer a versatile and bioinspired platform for wound healing and tissue regeneration, as they can self-assemble into nanoscale architectures that mimic the native extracellular matrix.^11–15^ By chemically modifying^16,17^ and co-assembling^18–20^ short peptides and amino acids, these scaffolds can promote cellular adhesion, while enabling the incorporation of antibacterial and regenerative bioactive motifs.^21–25^ We recently addressed the major limitations of CEAs by fabricating scaffolds that incorporate Fmoc-FRGD peptide into electrospun PCL. This scaffold supported dermal and epidermal layered organization and enhanced full-thickness skin regeneration without the need for additional cellular stratification, thereby improving both structural and functional outcomes.^26^ Recent advances in microfluidic-assisted 3D bioprinting further highlight how biofabrication technologies can be leveraged to incorporate bioactive components within complex scaffolds for regenerative applications.^27^

Despite these advances, most scaffolds are passive as they support the healing process but provide no means to track cellular behavior or regeneration dynamics over time. The absence of real-time physiological feedback is particularly critical in chronic wounds and complex tissue injuries.^28–30^ These challenges highlight the need for self-sensing scaffolds that not only guide tissue regeneration but also monitor healing processes *in situ* and in real-time.

Recent advances in wearable and implantable bioelectronics have yielded soft, flexible devices capable of sensing wound-relevant parameters such as temperature^31,32^, pH^33^, mechanical strain^34,35^, and tissue oxygenation.^36^ Among the various sensing modalities, electrical impedance spectroscopy (EIS) has emerged as a promising label-free technique for characterizing biological phenomena such as cell adhesion and proliferation.^37–39^

While these systems represent an important step toward digital wound care, they are typically designed as adjunctive electronics placed on the wound, rather than as integrated components of regenerative scaffolds. Their fabrication on non-porous, non-degradable substrates, often lacking conformability to dynamic wound geometries, further limits their long-term stability and biocompatibility.

Smart bandage technologies are particularly promising, enabling closed-loop wound care through the integration of multimodal sensors, electrical stimulators, and controlled therapeutic release.^40^ These platforms offer the potential for autonomous monitoring and treatment, reducing clinician intervention and improving patient outcomes. Nevertheless, most current devices rely on bulky, rigid electronic components and materials such as PEDOT:PSS or carbon nanotubes, which raise concerns regarding cytotoxicity, poor biodegradability, and limited tissue compatibility.^41–43^ Additionally, metal-based electrodes without proper adhesion to biological tissue often fail to maintain stable operation in the hydrated and mechanically dynamic wound environment.

To address these challenges, MXenes, a class of two-dimensional transition metal carbides and nitrides, have emerged as promising candidates for soft bioelectronics, owing to their high electrical conductivity, tunable surface chemistry, and biocompatibility.^44–46^ In particular, Ti_3_C_2_T_X_ MXene exhibits high capacitance and low cytotoxicity, making it suitable for applications in neural interfaces,^47,48^ wearable sensors,^49,50^ and tissue-integrated monitoring platforms.^51,52^ We previously demonstrated that a peptide-MXene hybrid hydrogel functions as a piezoresistive sensor, capable of transducing mechanical deformation into robust electrical signals while supporting cell viability.^53^ While these approaches each offer unique benefits, the real challenge lies in developing a single scaffold that enables continuous healing assessment while promoting full-thickness tissue regeneration.

Here, we present a multifunctional electrospun scaffold that addresses these unmet clinical and technological needs by seamlessly integrating regenerative bioactivity with real-time monitoring. We fabricated a scaffold composed of PCL nanofibers functionalized with Fmoc-FRGD peptides to promote cell adhesion and tissue regeneration. To integrate sensing functionality, biocompatible Ti_3_C_2_T_X_ MXene electrodes were conformally deposited onto the fibrous matrix, preserving the native architecture while enabling intimate electrical interfacing with the cells. This design supports label-free EIS measurements directly from the scaffold surface, allowing continuous assessment of cellular coverage and healing dynamics without disrupting the repair process (Figure 1a).

**Figure 1.**
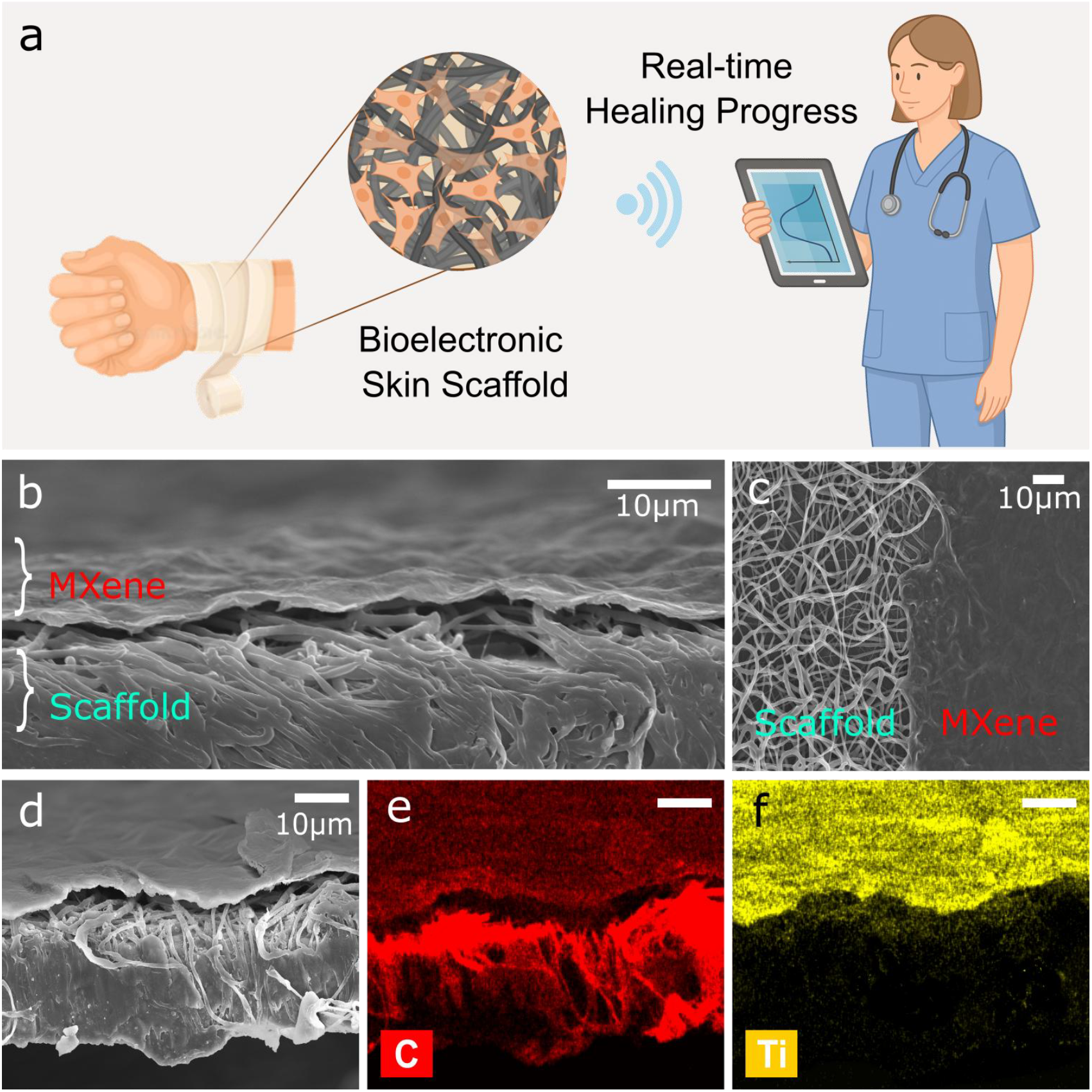
Fabrication and structural characterization of the bioelectronic skin scaffold. (a) Schematic illustration of the bioelectronic scaffold designed to support both regeneration and continuous monitoring of wound healing via impedance sensing. The scaffold enables real-time tracking of cellular proliferation and scaffold coverage, providing functional insights into the healing progression. (b-c) HR-SEM images showing (b) tilted and top-view morphology of the bioelectronic scaffold. (d-f) Element characterization of the scaffold using cross-sectional HR-SEM and EDS mapping. (d) Secondary electron image (e) Elemental mapping of carbon (red) (f) Elemental mapping of titanium (yellow). Scale bar = 10 µm.

## Results and Discussion

### Fabrication and Structural Characterization of the Bioelectronic Skin Scaffold

The scaffold was fabricated using electrospinning nonwoven nanofiber meshes of PCL fibers with a 2.5% Fmoc-FRGD peptide-to-PCL ratios, as previously reported.^26^ In a typical procedure, the components were mixed, dissolved, and dispensed through a single needle to obtain single-component fibers. To fabricate the sensing interface, an aqueous dispersion of Ti_3_C_2_T_X_ MXene was drop-cast onto two semicircular regions on the top surface of a 13 mm-diameter scaffold, one on the right and one on the left, forming opposing electrodes separated by a central gap. This configuration enables direct electrical examination of the regenerating tissue while preserving a biologically active region for cell growth.

Current-voltage (I-V) measurements were performed on the MXene electrodes patterned on the scaffold to assess their electrical performance. MXene-coated regions exhibited a linear, symmetrical response, consistent with Ohmic behavior and a low resistance of ∼ 70 Ω. In contrast, uncoated regions were electrically insulated (R ∼ GΩ) across the same voltage range (Figure S1). These results validate the formation of a continuous, conductive MXene interface capable of supporting stable electrical readout.

High-resolution scanning electron microscopy (HR-SEM) (Figure 1b) revealed a well-defined bilayer structure, with a conformal MXene film coating the upper surface of the fibrous scaffold. Top-view HR-SEM imaging (Figure 1c) showed a sharply defined boundary between the conductive and non-conductive regions, confirming the spatial precision of the deposition process. Cross-sectional SEM imaging further revealed that the MXene electrode formed a surface-confined layer with an average thickness of 10.4 ± 1.9 µm, overlaying a fibrous electrospun matrix 37.3 ± 3.5 µm in thickness (Figure S2).

To validate the elemental composition and spatial confinement of the conductive layer, we performed energy-dispersive X-ray spectroscopy analysis (EDS; Figure 1d-f). Carbon mapping (Figure 1e) showed a broad distribution across both scaffold and MXene-coated areas, while titanium mapping (Figure 1f), indicative of the Ti_3_C_2_T_X_ composition, was restricted to the upper layer. This spatial localization confirms that the MXene film remains surface-bound and does not infiltrate the scaffold’s interior, preserving its porous architecture and biological functionality.

### Evaluation of Cell Viability and Cell-scaffold Interactions

To evaluate the biocompatibility of the bioelectronic skin scaffold, we cultured NIH-3T3 fibroblasts on MXene-coated PCL-Fmoc-FRGD scaffolds and assessed cell viability and morphology over time (Figure 2). We first explored the influence of initial seeding quantity on viability across a 7-day period using the Alamar blue assay (Figure 2a). Cells were seeded at four quantities: 5k, 25k, 50k, and 500k cells per scaffold, and viability was quantified on days 1, 3, and 7. Across all conditions, cell viability was maintained or increased over time, particularly at 5k, 25k, and 50k, confirming that the MXene-electrode coated scaffold is biocompatible and supports cell growth. This is in correlation with the previous report on the biocompatibility of the Ti_3_C_2_T_X_ MXene.^54^

**Figure 2.**
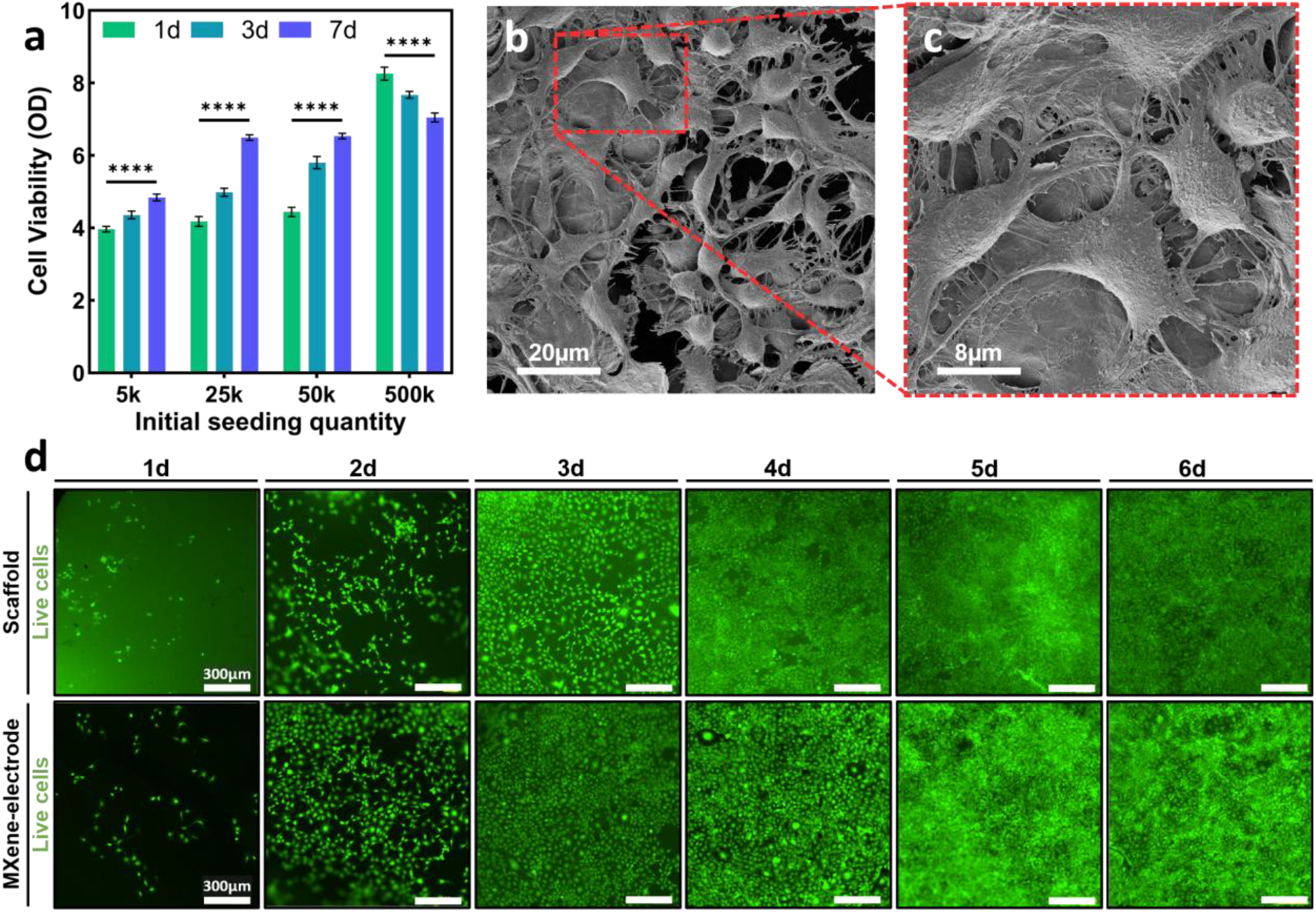
Evaluation of cell viability and spatial distribution on the bioelectronic scaffold. (a) Quantitative analysis of cell viability over time (days 1, 3, and 7) at varying initial seeding quantities (5k, 25k, 50k, and 500k cells/scaffold). (b-c) HR-SEM images of NIH-3T3 fibroblasts cultured on the scaffold (50k cells) after 7 days, showing extensive spreading, tight adhesion, and multilayered growth, indicating strong scaffold-cell interactions. Representative fluorescence images of live cells cultured on scaffold regions (top) and MXene-electrode regions (bottom) over 6 days (left to right). Scale bar = 300 µm.

Statistical analysis showed a significant increase in viability over time at low and intermediate seeding quantities (5k–50k), while the 500k group exhibited a slight, but significant, decline between days 1 and 7 (p < 0.0001). This reduction is likely due to excessive cell confluence within the confined culture area of the scaffold and the limited diffusion capacity of the 24-well. In contrast, the 50k condition maintained high and stable viability throughout the entire 7-day period, suggesting that it offers an optimal balance between cell quantity and environmental support. Additionally, it aligns with manufacturer recommendations for standard 24 well-plate culture protocols, supporting reproducibility and methodological consistency.^55^ Based on these results, we selected to continue with 50k cells per scaffold for all subsequent experiments.

HR-SEM imaging (Figure 2b-c) further demonstrated the scaffold’s ability to support robust cell adhesion, spreading, and spatial organization. Fibroblasts displayed an elongated morphology with prominent filopodia extending along and across fibers, indicating active engagement with the fibrous matrix. Cells were frequently observed bridging across pores and interacting with neighboring fibers, reflecting their response to the scaffold’s structural and topographical cues. In several regions, cells formed a multilayered structure (Figure 2b, S3), suggesting that the scaffold not only promotes initial attachment but also supports vertical growth and dense coverage over time.

To evaluate the spatial dynamics of cells across the scaffold, we performed fluorescence imaging of live cells over a 6-day period, comparing the scaffold’s non-coated area with the MXene-coated electrode regions (Figure 2d). Both regions demonstrated progressive increases in live cell coverage over time. By days 4-6, the cell coverage appeared to exceed 80% confluence across the scaffold surface, indicating that the MXene electrodes do not impede cell adhesion or proliferation, and support uniform coverage comparable to that of the non-coated fibrous matrix.

Taken together, these results establish that the MXene-coated PCL-Fmoc-FRGD scaffold provides a favourable microenvironment for fibroblast adhesion, proliferation, and survival over time.

### EIS Correspondence to Circuit Modelling

To evaluate the real-time sensing capability of the bioelectronic skin scaffold, we performed EIS analysis over a 9-day culture period. Impedance spectra were recorded every 2 hours across a frequency range of 0.1 Hz to 1 MHz for scaffolds seeded with NIH-3T3 fibroblasts (50k cells/scaffold) and for cell-free controls (Figure S4). To interpret the impedance behavior of the bioelectronic skin scaffold, we applied a modified Randles equivalent circuit model (Figure 3), a well-established framework for describing the electrochemical behavior at the scaffold-electrolyte interface, particularly under low-frequency regimes where interfacial processes dominate.^56^ This model effectively describes the key electrical features of our system, where minimal faradaic (redox) reactions occur and the impedance response arises primarily from capacitive and resistive components.

**Figure 3.**
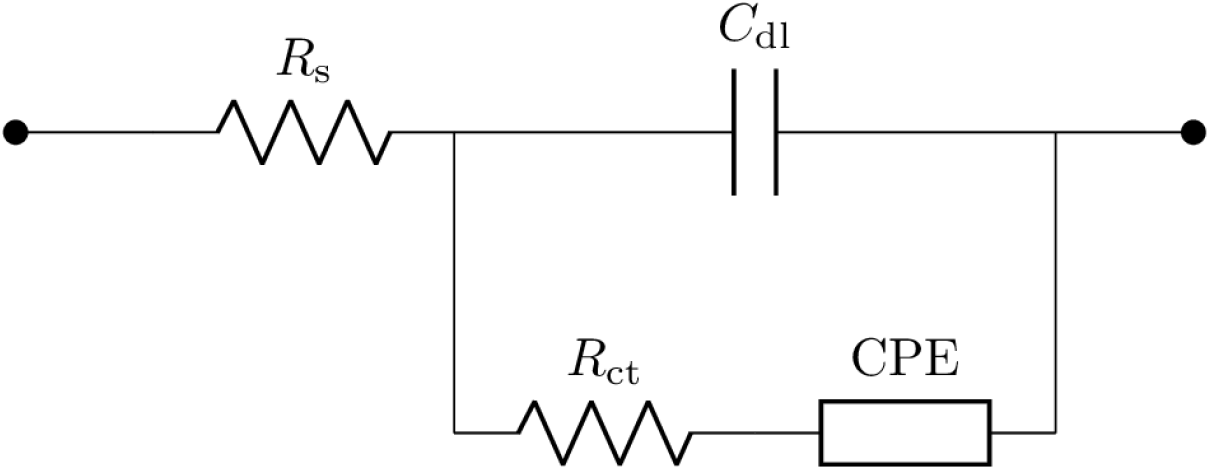
Equivalent electrical circuit model for impedance analysis of the bioelectronic skin scaffold. Schematic representation of the modified Randles circuit used to fit impedance spectra collected from the scaffold. The model includes solution resistance (Rs), charge transfer resistance (Rct), non-ideal double-layer capacitance (C_dl_), and a constant phase element (CPE) that captures frequency dispersion and ionic mass transfer effects at the scaffold–electrolyte–cell interface.

Specifically, Rs represents the series resistance of the electrolyte and electrodes, while charge transfer resistance (Rct) and the constant phase element (CPE) reflect the interfacial resistance and non-ideal capacitive behavior at the cell-scaffold interface, which may arise from factors such as surface roughness, cellular adhesion and coverage, or mass transport limitations. The double-layer capacitance C_dl_, modeled as a non-ideal or “leaky” capacitor, accounts for charge accumulation at the electrode-electrolyte interface and reflects changes in cellular adhesion and morphology over time. Given that the low-frequency regime is dominated by interfacial impedance, the Randles model is particularly well-suited for capturing dynamic changes in cell-scaffold interactions.

### Real-time Monitoring of the Bioelectronic Scaffold via Impedance Spectroscopy

The resulting data of the impedance spectra were fitted to the Randles equivalent circuit model, enabling the extraction of key interfacial parameters that reflect cell-scaffold interactions and dynamic changes in cellular coverage.

Bode plots of the impedance magnitude (|Z|) and phase angle (∠Z) revealed distinct behaviors between experimental conditions. In the cell-free scaffolds (Figure 4a), the impedance spectrum remained largely stable over time, reflecting a static interface dominated by the intrinsic conductivity of the MXene layer and the resistance of the surrounding electrolyte. A slight temporal drift was observed, which may be attributed to a combination of gradual surface oxidation of the MXene, potentially reducing its conductivity^57^ and nonspecific adsorption of serum proteins or ions from the culture medium, altering the interfacial impedance.^58^

**Figure 4.**
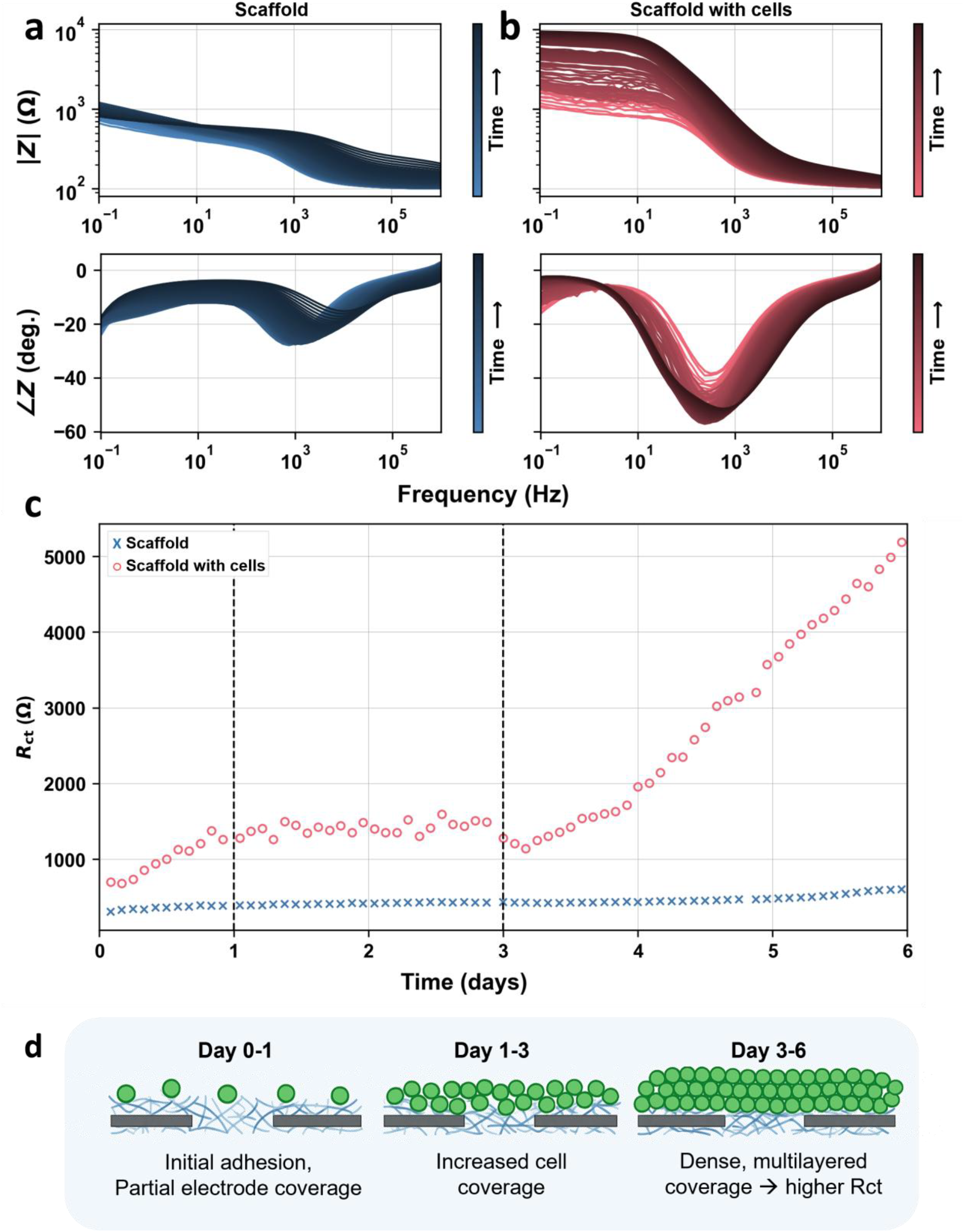
Real-time impedance monitoring of cell growth on the bioelectronic scaffold. (a-b) Bode plots showing impedance magnitude (|Z|, top row) and phase angle (∠Z, bottom row) as a function of frequency over time, for (a) scaffolds without cells (left, blue) and (b) Scaffolds seeded with 50,000 cells/scaffold (right, red). Arrows and color gradients indicate the progression of time. (c) Rct values over time extracted from equivalent circuit fitting of impedance spectra, comparing cell-free (blue) and cell-seeded (red) scaffolds. (d) Schematic illustration correlating impedance dynamics with cellular behavior. Three biological stages were observed: initial adhesion and partial coverage (Days 0-1), spreading and monolayer formation (Days 1-3), and dense, multilayered growth (Days 3-6).

In contrast, cell-seeded scaffolds exhibited progressive shifts in both magnitude and phase (Figure 4b, color gradient), particularly at low frequencies (<100 Hz), where we observed a clear increase in impedance magnitude, indicative of evolving interfacial dynamics driven by cell adhesion and interaction with the scaffold matrix.

Fitting the spectra to the Randles circuit enabled the extraction of the charge transfer resistance (Rct), a sensitive parameter that reflects cell-electrode interfacial processes, such as the proliferation and coverage of cells on the electrode surface. This parameter becomes more prominent at lower frequencies, where interfacial phenomena dominate the impedance response. As shown in Figure 4c, Rct remained constant in the cell-free scaffolds (∼500-600 Ω) but increased significantly over time in the presence of cells, rising from ∼800 Ω on day 0 to over 5000 Ω by day 6.

The changes of Rct over time revealed distinct stages corresponding to key stages of cell growth and scaffold coverage. During the initial 24 hours (Day 0-1), the Rct increased rapidly from ∼370 Ω to ∼1,280 Ω, representing a ∼3.4-fold rise. This rapid increase corresponds to early cell adhesion and partial scaffold coverage, during which the interface became increasingly obstructed by attached cells, thereby impeding the flow of charge carriers across the scaffold and electrolyte. From Day 1 to Day 3, the Rct plateaued within the range of ∼1,300-1,600 Ω, reflecting a stabilization in monolayer formation and moderate changes in cell morphology. This stage likely corresponds to cell spreading, matrix remodelling, and initial stratification over the conductive surface. Between Day 3 and Day 6, a more pronounced elevation in the Rct was observed, rising from ∼1,950 Ω to ∼5,500 Ω, a >15-fold increase relative to the baseline. This sharp rise is consistent with dense multilayered cell growth that substantially increased the barrier to charge transfer, likely due to enhanced cell-scaffold interactions and extracellular matrix deposition at the scaffold interface. Such interpretation is supported by HR-SEM images, showing vertically stacked fibroblasts and extended filopodia (Figure 2b-c, Figure S3), as well as fluorescence staining indicating uniform high viability and cells’ confluence (Figure 2d). These findings are further illustrated by the mechanistic schematic in Figure 4d, which depicts how increasing cell density progressively impedes ion transfer at the scaffold-electrolyte interface. As cells adhere and proliferate, their membranes, being electrically insulating by nature, act as barriers to current flow, thereby increasing the Rct in proportion to cell coverage. This dynamic behavior demonstrates the scaffold’s ability to track key cellular processes, including adhesion, proliferation, and multilayer formation, in a label-free manner. Accordingly, Rct serves as a sensitive indicator of cell growth over time.

### Correlation Between Standard Cell Viability Assays and Impedance Sensing

To validate the biological relevance of the impedance-based sensing platform, we systematically correlated Rct values with two gold-standard biological assays: metabolic cell viability and fluorescence-based cell staining based coverage. As shown in Figure 5a, NIH-3T3 fibroblasts cultured on the bioelectronic scaffold exhibited a progressive increase in metabolic activity over a 9-day period, as measured by the Alamar Blue assay. Correlating this biological trend with Rct values, extracted from the real-time impedance spectra, revealed a strong linear relationship, with a high degree of accuracy (R^2^ = 0.9678) (Figure 5b).

**Figure 5.**
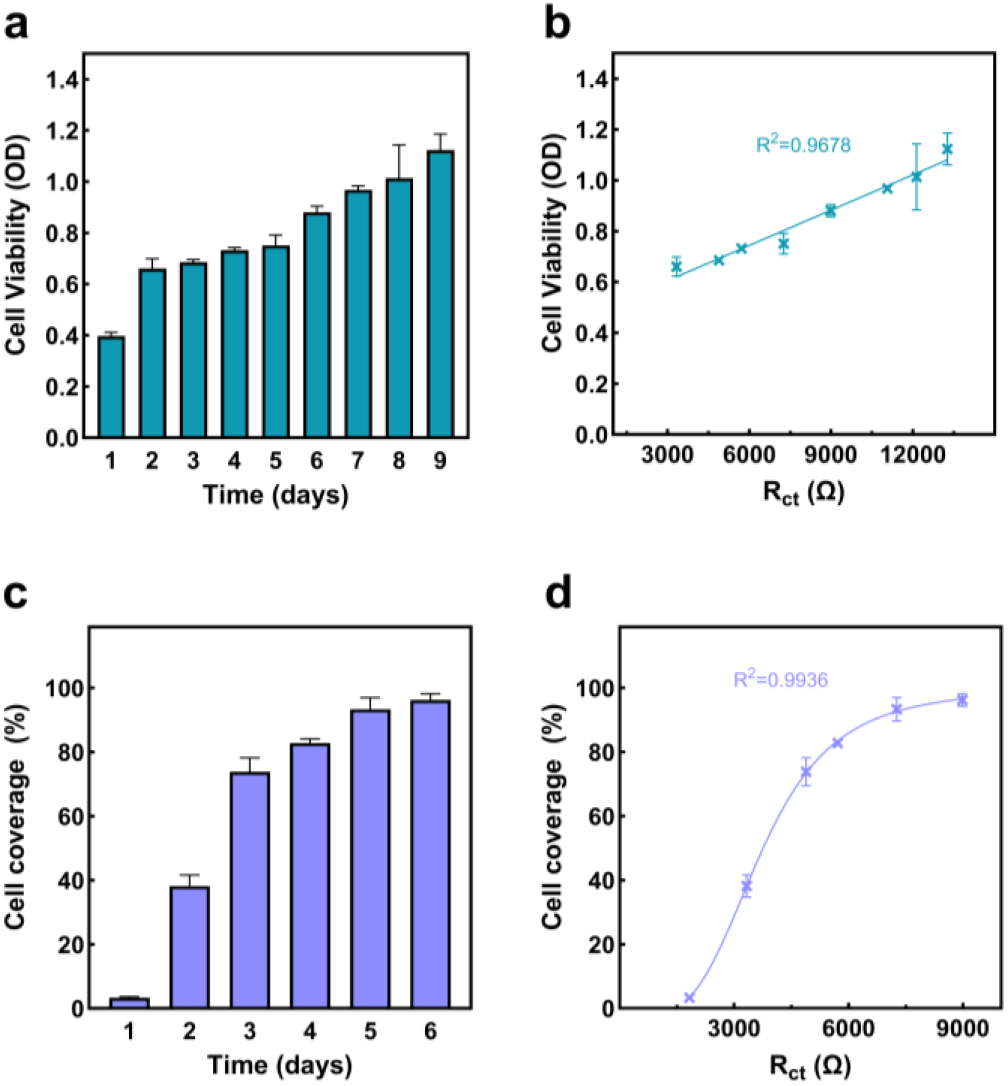
Correlating cell viability and coverage assays with real-time impedance measurements. (a) Cell viability by Alamar Blue assay showing a time-dependent increase in metabolic activity of NIH-3T3 cells cultured on the bioelectronic scaffold. (b) Linear correlation between cell viability and Rct extracted from impedance spectra (R^2^=0.9678). (c) Quantification of cell coverage over time based on live cell fluorescence staining. (d) Sigmoidal correlation between cell coverage and Rct values, modeled using a 4-parameter logistic function (R^2^=0.9936).

To complement the metabolic viability analysis, we tracked cell coverage dynamics using fluorescence-based staining of live cells (Figure 5c). Confluence increased sharply during the first three days, then reached a plateau from day 4 through day 6, indicating maximal surface occupancy. When plotted against the corresponding Rct values (Figure 5d), the correlation exhibited a classic sigmoidal profile that was exquisitely captured by a four-parameter logistic (4PL) regression (R^2^ = 0.9936).

Together, these results demonstrate that Rct serves as a sensitive, real-time indicator of both cellular viability and surface coverage, offering quantitative resolution without the need for disruptive labeling or endpoint assays. The close convergence of optical and electrical readouts further supports the utility of our approach for continuous monitoring in wound closure assessment.

### Conclusions

In this study, we developed a multifunctional bioelectronic skin scaffold that seamlessly integrates regenerative and diagnostic functions within a single platform. For that aim, we utilize the previously validated scaffold of electrospun PCL functionalized with the bioactive peptide Fmoc-FRGD, which has been shown to support full-thickness skin regeneration. We then introduced a surface-confined MXene (Ti_3_C_2_T_x_) layer to endow the scaffold with electrical conductivity and real-time sensing capabilities. Structural and morphological characterization confirmed the bilayer architecture and the stable confinement of the MXene to the scaffold surface, preserving its biological performance.

EIS was used to continuously monitor cell behavior over a 9-day period. Impedance spectra fitted to a Randles equivalent circuit model revealed that the Rct parameter serves as a sensitive, label-free, and non-destructive indicator for cell adhesion, proliferation, and multilayer formation. Temporal changes in Rct were tightly correlated with standard biological assays, including Alamar Blue viability measurements and fluorescence imaging, confirming the scaffold’s ability to quantitatively resolve key biological events. Moreover, the scaffold maintained high cytocompatibility and supported robust, multilayered fibroblast coverage.

This convergence of regenerative support with electrical functionality positions the MXene-based scaffold as a next-generation platform for real-time wound sensing. Its ability to continuously monitor cell proliferation, implying the healing progress without disrupting the wound site, offers a potentially powerful solution for managing chronic wounds, including diabetic ulcers, pressure sores, and burns. Beyond wound care, the technology may be extended to other applications in tissue engineering, implantable diagnostics, and smart regenerative systems, where regeneration and real-time feedback is essential for therapeutic success.

### Experimental Section

#### Preparation of the bioelectronic scaffold

For the electrospinning of PCL fibers (average Mn 80,000, Sigma-Aldrich (Rehovot, Israel)) containing Fmoc-FRGD (GL Biochem (Shanghai, China)), 0.450 g of PCL was first dissolved in 2.775 mL of hexafluoroisopropanol (HFIP) (Sigma-Aldrich (Rehovot, Israel)), to obtain a clear homogeneous solution. Separately, Fmoc-FRGD was dissolved in HFIP at a concentration of 50 mg/ml. Next, the Fmoc-FRGD stock solution was added to the PCL solution to obtain a final composition of 15% (w/v) PCL and 0.375% (w/v) Fmoc-FRGD, corresponding to a peptide-to-PCL ratio of 1:40 (2.5%). Fresh peptide stock solutions were prepared for each experiment, and the final mixed solution was used immediately.

The electrospinning setup contained a syringe pump (New Era Pump Systems Inc. (NY, USA)), a power supply (DC voltage source, Gamma High Voltage Research (Florida, USA)), a moving stage, and a rotating drum collector. The polymeric solutions were dispensed via a 25-gauge needle at a constant flow rate of 0.450 mL/hr. A driving voltage of 7 kV resulted in a stable jet, and the non-woven fiber meshes were collected on a rotating drum covered with nonstick aluminum foil, onto which circular cover glasses 13mm in diameter were placed. The tip-to-ground distance was set to 7 cm, and the drum rotating speed was set to 80 rpm.

#### Ti_3_AlC_2_ MAX and Ti_3_C_2_T_z_ MXene synthesis

Ti_3_C_2_T_z_ MXene was synthesized by selectively etching the metallic Al layer of the Ti_3_AlC_2_ MAX precursor. The Ti_3_AlC_2_ MAX precursor was synthesized by mixing titanium carbide (Alfa Aesar, 99.5%, 2 µm), aluminum (Alfa Aesar, 99.5%, 325 mesh), and titanium (Alfa Aesar, 99.5%, 325 mesh) powders at a 2:1.15:1 molar ratio, respectively. The elementary powders were mixed in a tumbler mixer at 150 rpm for 24 h until a homogeneous mixture was obtained. The mixed powders were then heat-treated in a tube furnace at 1400 °C at a heating rate of 5 °C min^−1^ for 3 h under a flowing Ar atmosphere (100 sccm). Finally, the resulting Ti_3_AlC_2_ MAX phase was milled to a fine powder using a planetary ball mill and sieved through a 400-mesh screen (<38 µm) to attain a homogeneous powder.

The Ti_3_AlC_2_ powder was etched in an HCl and LiF solution, which reacted to form in situ HF. Initially, 1.6 g of LiF (Alfa Aesar, 99.5%, 325 mesh) was dissolved in 20 mL of 32% HCl (Fisher Scientific). This was followed by the slow addition of 1 g of the Ti_3_AlC_2_ powder to prevent a violent reaction, and the solution was then stirred for 24 h at 45 °C and 300 rpm. The etching was performed in an HDPE container with a vent to allow the release of hydrogen produced during the reaction. The black slurry was transferred to a 50-mL centrifuge vial, and type I deionized (DI) water was added to fill the vial. The slurry was centrifuged at 2300 RCF for 2 min, and the resulting clear supernatant was decanted while retaining the bottom sediment. To remove the excess LiF, the etched material was washed twice with 1M HCl solution and centrifuged after each wash. After the HCl washing, the material was washed and centrifuged with the above DI water several more times, until the pH of the solution reached ∼ 6. This step yielded a multilayered Ti_3_C_2_T_X_ MXene. To exfoliate the MXene to a single layer of MXene, the mixture was sonicated under a bubbling Ar flow for 1h, while maintaining the temperature of the sonication bath below 10 °C. The solution was then centrifuged for 30 min at 3500 RPM, and only the colloid solution was collected.

#### Integration of MXene electrodes into electrospun scaffolds

To fabricate MXene electrodes on the electrospun scaffold, a 6 mm-wide hydrophobic silicone spacer was centrally positioned on the scaffold surface to mask the middle region. MXene solution (10 mg/mL in deionized water) was then drop-cast (18 µL per side) onto the exposed regions flanking the spacer. The scaffolds were placed in a desiccator and allowed to dry overnight to ensure complete solvent evaporation and MXene film formation. After drying, the silicone spacer was carefully removed, yielding two well-defined MXene electrode pads separated by a 6 mm gap.

#### Cell viability assay

MC3T3-E1 mouse fibroblast cells were purchased from ATCC and cultured in Dulbecco’s Modified Eagle’s Medium (DMEM)(Gibco) supplemented with 10 wt% fetal bovine serum (FBS), 1% Penicillin-Streptomycin (Sigma-Aldrich), and 1% L-Glutamine (200 mM) (Sartorius). The cells were cultured in a petri dish at 37 °C in a humidified atmosphere containing 5% CO_2_. The bioelectronic scaffolds were seeded at a concentration of 50,000 cells/scaffold with DMEM medium and were incubated for 24 hours to allow cell attachment. Cell viability was evaluated using the AlamarBlue™ reagent (ThermoFisher). The cell culture medium was replaced with AlamarBlue diluted 1:9 in supplemented DMEM and incubated for 4 h (37 °C, 5% CO_2_). Then, the AlamarBlue/DMEM solutions were transferred to a clean well plate and absorbance was measured using a Tecan Spark Infinite M200PRO plate reader at λ = 570 nm and λ = 600 nm. After the AlamarBlue analysis, the scaffolds were washed once with PBS to remove AlamarBlue residues, and fresh DMEM medium was then added. This process was repeated every 24 hours for 9 days.

#### High-resolution scanning electron microscopy (HR-SEM)

HR-SEM was performed in a Zeiss Gemini 300, in a high vacuum mode, at a work distance (WD) ≈ 10 mm and voltage of 3 kV. After electrospinning and MXene depositing, the scaffolds were coated with Au using a sputtering system. Energy dispersive spectrometer (EDS) was used for elemental composition mapping of the scaffold. HRSEM images were analyzed using ImageJ software to determine the layers average thickness (n = 30). To visualize cell attachment to the scaffold, scaffolds were seeded with 50,000 cells each in DMEM medium and incubated for 7 days. Following incubation, the scaffolds were washed three times with PBS and fixed overnight at 4 °C using 2.5% (v/v) glutaraldehyde in PBS. Post-fixation, samples were washed once with PBS and subjected to a graded ethanol dehydration series (50%, 75%, and 100% in PBS), with each step lasting 30 minutes. Subsequently, the samples were maintained in 100% ethanol before undergoing critical point drying, gold sputter-coating, and HR-SEM imaging.

#### Live cell staining

Fluorescent Live staining assay (Sigma Aldrich) containing fluorescein diacetate (6.6 μg/ml) was used to visualize the proportion of viable cells. The labelled cells were incubated for 5 minutes (37 °C, 5% CO_2_), rinsed once with PBS, and then immediately viewed using a confocal ZEISS LSM 900 microscope (ZEISS, Germany).

#### Real-time electrical impedance spectroscopy (EIS)

Cell-seeded scaffolds were placed in a 24-well cell repellent plate containing the above-mentioned supplemented DMEM. A custom-designed lid for the 24-well plate was fabricated, featuring two electrode ports per well spaced 1 cm apart. Two platinum wires were inserted into each pair of ports, making direct contact with the MXene-coated electrodes on the scaffold surface. A multi-channel impedance analyzer (MultiPalmSens 4, PalmSens Inc.) was connected to the platinum electrodes to perform real-time EIS measurements. Impedance spectra were recorded every 2 hours over a period of 6 days, across a frequency range of 0.1 Hz to 1 MHz using a sinusoidal voltage amplitude of 100 mV and a DC offset of 0 V. Each condition was tested in triplicate, with an unseeded scaffold serving as a negative control. All measurements were conducted inside a standard incubator (37 °C, 5% CO_2_) to maintain physiological conditions.

#### Electrical modelling

Impedance spectra were acquired from cell-seeded scaffolds and were fitted to an equivalent circuit model (Randles model with a leaky capacitor), using a parametrized form of the circuit’s analytical impedance characteristics. The model was then fitted to the experimental measurements using dual annealing global optimization,^59^ as implemented in SciPy,^60^ to obtain best-fit values of the equivalent circuit parameters. The accuracy of the fitting was quantified using the mean absolute error of the fit, in the complex domain. This error was observed to be less than 7% for all experimental measurements, indicating proper fitting.

#### Correlation of cell viability with Rct by linear regression

Cell viability (OD) was determined by the Alamar Blue assay at eight distinct time points (n = 1 per time point; total n = 8). The earliest timepoint (day 1), corresponding to early adhesion prior to metabolic stabilization, was assessed separately. Its inclusion in the correlation analysis produced only a negligible change in R^2^ (Figure S5). In parallel, Rct was extracted from real-time electrochemical impedance spectra. OD Viability values were plotted against Rct (Ω), and a simple linear regression was performed in GraphPad Prism 10.4.2 using ordinary least-squares fitting. The regression equation was:

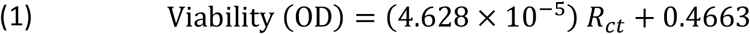

With coefficient of determination R^2^ = 0.9678. Goodness‐of‐fit statistics confirming a highly significant, positive correlation between Rct and metabolic viability.

### Correlation of cell coverage with Rct by four-parameter logistic regression

Cell coverage (%) was quantified from fluorescence live cell images acquired at six time points (days 0-6). In parallel, Rct was extracted from real‐time electrochemical impedance spectra. These data were plotted in GraphPad Prism 10.4.2 and fit to a four-parameter logistic (4PL) model according to the equation:

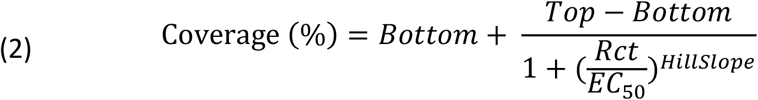

The best-fit parameters (±95% CI) were:

- Bottom = -3.132 (-8.349 to 0.929)
- Top = 99.91 (96.82 to 103.7)
- EC_50_ = 3702 (3570 to 3828)
- HillSlope = 3.825 (3.280 to 4.379)

Goodness-of-fit metrics were R^2^ = 0.9936, confirming an excellent sigmoidal correlation between scaffold coverage and Rct.

#### Statistical analysis

All statistical analyses were performed using Prism version 10.4.2 (GraphPad). Data is presented as mean ± standard deviation (SD), based on at least three independent experiments. Details of specific statistical tests and comparisons are provided in the respective figure legends.

## Acknowledgments

This work was supported by the European Research Council (ERC), under the European Union’s Horizon 2020 research and innovation program (grant agreement no. 948 102) (L.A.-A.). This work was supported by the Israel Science Foundation (ISF grant: 2422/24) (L.A.-A.). This work was supported by the Israel Science Foundation (ISF grant: 2527/22) (M.S.). The authors acknowledge the support of the Zimin Institute (L.A.-A. and M.S). D.C.-G. acknowledges the Marian Gertner Institute for Medical Nano systems at Tel Aviv University. D.C.-G. gratefully acknowledges the support of the Colton Foundation. The authors acknowledge the Chaoul Center for Nanoscale Systems of Tel Aviv University for the use of instruments and staff assistance. The authors thank the members of the Adler-Abramovich, Shacham-Diamand, and Sokol groups for the helpful discussions.

