## Supplementary Information for "A Bioelectronic Scaffold for Label-Free, Real-Time Monitoring of Wound Healing"

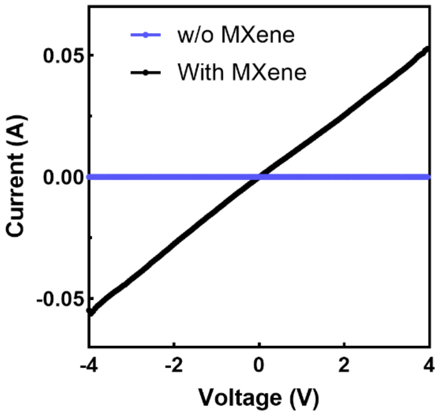


**Figure S1. Electrical performance.** I–V characteristic curves of the electrospun scaffold with (black) and without (blue) a MXene $Ti_3C_2T_x$ layer.

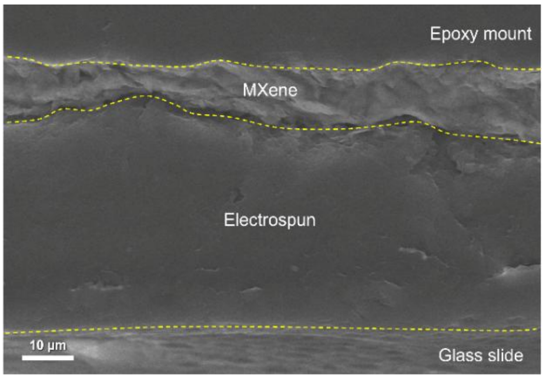


**Figure S2. Morphological characterization of the bioelectronic scaffold.** Cross-sectional SEM image of the MXene-coated scaffold, showing the layered architecture following epoxy embedding and microtoming. The $Ti_3C_2T_x$ MXene layer (highlighted between yellow dashed lines) forms a conformal, surface-confined coating above the electrospun PCL/Fmoc-FRGD scaffold. The clear interface and uniform thickness demonstrate that the conductive film remains localized at the surface, preserving the porous internal structure of the scaffold. Scale bar = 10 µm.

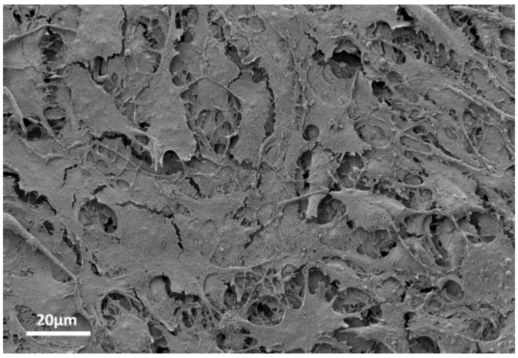


**Figure S3. HR-SEM image showing fibroblasts cultured on the MXene-coated PCL-Fmoc-FRGD scaffold.** Cells exhibit multilayered organization and elongated morphology, with filopodia extending across fibers and between cell layers, suggesting advanced colonization and stable scaffold integration. Scale bar = 20 µm.

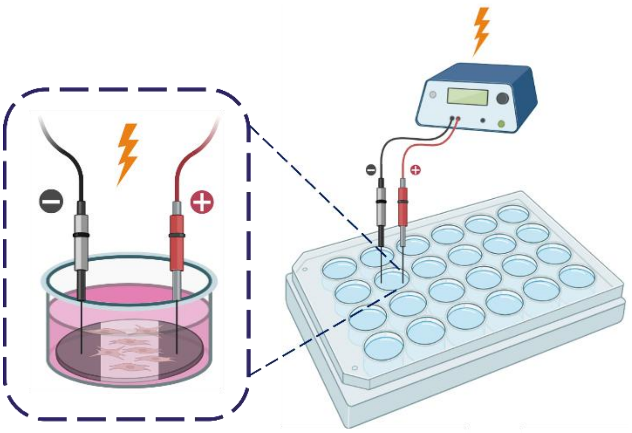

**Figure S4. Schematic illustration of the experimental setup for impedance measurements.** Cell-seeded scaffolds were placed in a 24-well plate, with MXene-coated electrodes connected to a potentiostat via platinum wires. Electrical signals were applied across the scaffold to enable real-time impedance monitoring of cell behavior under physiological culture conditions.

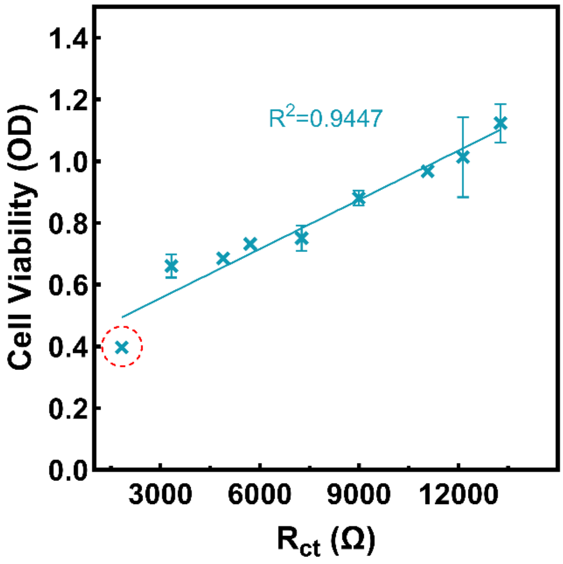


**Figure S5. Impact of Day 1 data point on the correlation between Rct and cell viability.** The full dataset including the day 1 timepoint is shown. The early timepoint exhibits lower metabolic activity and Rct values that slightly deviate from the overall trend, likely due to incomplete cell adhesion and minimal metabolic engagement. As a result, this point was excluded from the main analysis (Figure 5b) to better reflect the stable biological correlation.
